# Cortical tracking of natural speech by children with developmental language disorder (DLD): An EEG speech decoding investigation

**DOI:** 10.1101/2025.07.14.664754

**Authors:** Mahmoud Keshavarzi, Susan Richards, Georgia Feltham, Lyla Parvez, Usha Goswami

## Abstract

The sensory/neural Temporal Sampling (TS) theory of developmental language disorder (DLD) is based on the sensory and linguistic impairments in rhythm processing that are found in children with both developmental dyslexia (DD) and DLD. These sensory/linguistic impairments include decreased sensitivity to amplitude rise times (ARTs, the sensory triggers related to automatic cortical speech tracking), syllable stress patterns and speech rhythm. At the neural level TS theory predicts impairments in the cortical tracking of different rates of amplitude modulation (AM) in the speech signal <10Hz. To date, the accuracy of low-frequency cortical tracking in natural continuous speech has not been measured in children with DLD. Here, EEG was recorded during story listening from children with and without DLD aged around 9 years, and decoding analyses in the delta, theta and alpha (control) bands were carried out. EEG power was computed in the delta, theta and gamma bands, and phase-amplitude coupling and phase-phase coupling (PAC, PPC) were also computed between bands. Whole-brain analyses showed that the accuracy of low-frequency decoding (delta, theta) did not differ between groups. However, region-specific analyses revealed significantly reduced delta-band speech tracking in the right temporal cortex in the children with DLD. PAC and PPC dynamics did not differ between groups. The data suggest that low-frequency cortical tracking impairments in DLD may be spatially constrained to the right hemisphere rather than globally present as in DD. The data are discussed using TS theory.

**Highlights:**

- Cortical tracking of natural speech is measured in children with and without DLD
- Delta-band tracking is impaired in right temporal regions in children with DLD
- Cross-frequency coupling dynamics (PAC, PPC) did not differ between groups

## 1. Introduction

The potential sensory/neural causes of developmental language disorder (DLD) are poorly understood. Children with DLD show language difficulties that cause functional impairments in many aspects of linguistic processing, yet to date there is no agreed aetiology. For example, the CATALISE multi-national study of DLD identified linguistic processing problems with syntax, such as distinguishing grammatical from ungrammatical sentence forms, problems with word semantics, such as showing a limited understanding of word meanings, impairments in verbal short-term memory, and difficulties with phonology (Bishop et al., 2017). Given this wide range of impairments, the underlying aetiology seems likely to affect fundamental sensory/neural aspects of processing the speech signal. One prior avenue of interest has been whether hemispheric asymmetries during linguistic processing may be important in aetiology (Bishop, 2013). Using fMRI, it has been shown that left frontal and temporal cortical areas show reduced activation in DLD children during language processing (Asaridou & Watkins, 2022, for review). However, the fMRI findings do not demonstrate which speech processing mechanisms may be affected, and indeed laterality differences per se do not seem as fundamental as once thought (Parker et al., 2022). Prior MEG studies of DLD using speech processing tasks are consistent with the fMRI literature in that they demonstrate atypical left hemisphere function, particularly in temporal regions (Helenius et al., 2009, 2014). Again, however, these MEG demonstrations have not provided mechanistic information about the underlying neurobiological basis of DLD.

By contrast, the Temporal Sampling (TS) theory of childhood language disorders, originally proposed on the basis of experimental data from children with developmental dyslexia (DD, Goswami, 2011), does enable predictions concerning aetiology. TS theory builds on recent mechanistic advances in the neuroscience of speech processing, specifically the oscillatory framework proposed by Giraud and Poeppel (2012), which offers new opportunities for understanding developmental language disorders like DD and DLD (Goswami, 2011, 2015, 2022). Regarding DLD, TS theory utilises acoustic parameters of the speech signal related to processing speech rhythm and prosody, such as amplitude (loudness) and frequency (pitch) modulation (AM, FM). AM and FM changes in the continuous speech signal characterise all languages, and the processing of rhythm has been shown to be impaired in children with DLD in a range of languages (Corriveau et al., 2007; Corriveau & Goswami, 2009; Przybylski et al., 2013; Cumming et al., 2015; Richards & Goswami, 2015, 2019; Sallat & Jentschke, 2015; Bedoin et al., 2016). Indeed, sensitivity to speech rhythm is thought to be a cross-language precursor of language acquisition by infants (Mehler et al., 1988; Nazzi et al., 1998). Accordingly, difficulties with rhythm processing could be expected to have a profound effect on language acquisition. Furthermore, speech rhythm patterns in the continuous speech signal are cued by amplitude rise times (ARTs; Greenberg et al., 2003), and English-speaking children with DLD exhibit impaired discrimination of ART in non-speech tasks (Corriveau et al., 2007; Fraser et al., 2010; Beattie & Manis, 2012; Cumming et al., 2015; Richards & Goswami, 2015, 2019). This difficulty in discriminating ARTs could be expected to affect both speech rhythm perception and also the neural encoding of the continuous speech signal. ARTs (“speech edges”) are thought to phase-reset oscillatory networks in auditory cortex, thereby enabling accurate speech-brain synchronisation (Giraud & Poeppel, 2012; Doelling et al., 2014).

To explain the speech rhythm impairments that characterise children with DLD, TS theory has proposed that sensory/neural impairments in the cortical tracking of low-frequency AM rates < 10 Hz (known to carry information about prosody, see Goswami, 2011; Greenberg, 2006), already known to characterise children with DD (Goswami, 2022, for recent review), may also characterise children with DLD. A key focus of TS theory is the different bands of low-frequency AMs that are nested within the speech amplitude envelope (AE) of child-directed and infant-directed speech, particularly bands of AMs centred on ∼2 Hz and ∼5 Hz (corresponding to the EEG delta and theta bands, 0.5–4 Hz and 4–8 Hz respectively; see speech modelling studies by Leong & Goswami, 2015; Leong et al., 2017). These different AM bands underpin both rhythm perception and neural speech encoding across languages (Daikoku & Goswami, 2022, for review). Meanwhile, the speech modelling studies of infant- and child-directed speech (IDS, CDS) show that *phase relations* between these bands of AMs (bands centred on ∼2Hz and ∼5Hz) enable the parsing of phonological information at the prosodic and syllabic levels (‘acoustic-emergent phonology’, see Leong & Goswami, 2015; Leong et al., 2017). Accordingly, the accurate cortical tracking of delta- and theta-band speech information and phase-phase coupling (PPC) during infancy and early childhood may be particularly important in supporting the development of an emergent linguistic system (Leong et al., 2017). By TS theory, prosodic learning in all languages may be related to AM-based speech encoding mechanisms that are present from birth, with the accuracy of encoding lower frequencies <10 Hz of particular developmental importance (Goswami, 2022). Low-frequency speech-brain cortical tracking is already measurable in infancy (Kalashnikova et al., 2018; Jessen et al., 2019; Ortiz Barajas et al., 2021; Attaheri et al., 2022; Menn et al., 2022).

Regarding DLD therefore, by TS theory neural encoding of low-frequency speech envelope information would be expected to be impaired in children with DLD. To our knowledge, the fidelity of the cortical tracking of continuous speech by oscillatory neural networks has yet to be investigated in children with DLD. This is surprising, as studies of the cortical tracking of continuous speech by children with DD are frequent in the literature, using both EEG and MEG approaches (Power et al., 2016; Molinaro et al., 2016; Di Liberto et al., 2018; Destoky et al., 2020, 2022; Keshavarzi et al., 2022a; Mandke et al., 2022). In these DD studies, the cortical tracking of speech by children with dyslexia was selectively impaired in frequency bands below 10Hz, supporting TS theory. Affected children in English, French and Spanish all showed decreased speech-brain synchronisation in the delta (and sometimes theta) bands, in both sentence listening and story listening tasks. Furthermore, longitudinal TS-driven studies of infant language acquisition show that the accuracy of cortical tracking of continuous speech in the delta band measured at 11 months is a significant predictor of language outcomes at 2 years (Cambridge BabyRhythm study, Attaheri et al. 2024). This suggests a core role for the fidelity of delta-band cortical tracking in efficient linguistic development. The Cambridge BabyRhythm study included 120 TD infants, and also reported that the accuracy of theta-gamma phase-amplitude coupling (PAC) to continuous speech was a second predictor of enhanced language outcomes. Meanwhile, infants exhibiting higher theta power (power spectral density, PSD) and a higher theta/delta ratio during continuous speech listening had worse language outcomes (Attaheri et al., 2024). These prior TS-driven developmental data suggest that children with DLD might be expected to have poorer delta band cortical tracking of continuous speech, excessive theta power when listening to continuous speech, and worse theta-gamma PAC.

Oscillatory EEG and MEG studies with children with DLD are beginning to appear in the literature, but to date these studies have not utilised continuous speech stimuli such as sentences or stories. Instead, current studies rely on paradigms using single words or speech features within words (such as target phonemes), or even nonspeech stimuli or no task. Prior resting state studies may be indicative of oscillatory differences that are related to language development. For example, Lum et al. (2022) reported a negative correlation between resting state power in the theta band and TD children’s scores on the Clinical Evaluation of Language Fundamentals (CELF; Wiig et al., 2017) Recalling Sentences task, which is a diagnostic measure of DLD in English. No other associations between CELF scores and oscillatory power were found. Theta band resting state findings were also reported in a study with TD Chinese-speaking children, who were aged around 10 years (Meng et al., 2022). Meng et al. reported that resting state theta power was negatively associated with a vocabulary subtest of the Wechsler Intelligence Scale for Children (Vocabulary Definitions subtest). Accordingly, resting state studies are suggestive of enhanced oscillatory power in TD children with poorer language development. In an MEG study with DLD children aged 10–15 years, the first (to our knowledge) oscillatory study of word listening reported atypical oscillatory responses compared to age-matched typically-developing (TD) children (Nora et al., 2024). Nora et al. reported that early cortical decoding (100 ms latency) of individual words in Finnish did not differ between children with DLD and TD children, for either amplitude envelope or speech spectrogram measures. However, at longer latencies (at ∼200–300 ms lags) the data from the children with DLD showed poorer cortical representation of the AE information. This finding was interpreted by Nora et al. (2024) as reflecting poorer retention of acoustic-phonetic information in short-term memory, consistent with the verbal memory problems that characterise children with DLD across languages.

The oscillatory framework that motivated TS theory (Giraud & Poeppel, 2012) also identifies speech-brain synchronisation at faster rates such as beta (∼15–25 Hz) and gamma (> 30 Hz). In the Giraud and Poeppel framework, phonemic information in the speech signal is encoded neurally by gamma band responses (Lehongre et al., 2011, 2013). Phonemic representation in DLD is known to be impaired from nonword repetition studies, in which affected children make phoneme-level errors when repeating nonwords such as ‘loddernapish’ and ‘fenneriser’. Interestingly, *prospective* studies of infants at potential family risk for language disorders indicate that resting state power in the faster gamma band is associated with later DLD. These studies all used non-speech stimuli such as pairs of tones (Benasich et al., 2008; Choudhury & Benasich, 2011; Gou et al., 2011; Cantiani et al., 2016, 2019). Some of these infant studies also measured resting state theta power, which was not found to predict later language outcomes (Cantiani et al., 2016). Accordingly, the oscillatory infant and child data available to date, which are primarily resting state data, appear to identify at least two neurophysiological frequency bands as potentially atypical in DLD, the gamma band in infancy and the theta band in childhood. However, as none of these infant studies involved linguistic processing, these studies cannot provide a test of the TS framework.

One exception to the DLD field’s reliance on single word or nonspeech paradigms is a TS-driven oscillatory EEG study that used a story listening task (Araújo et al., 2024). Araújo et al. reported that children with DLD aged on average 9 years had significantly greater delta-theta PAC compared to TD children during natural speech listening. This may suggest that low-frequency oscillatory dynamics are atypical in the DLD brain during natural speech listening. However, only 7 children with DLD were tested in this study. A further TS-inspired EEG study conducted with the same 9-year-old children tested here measured neurophysiological responses to rhythmic speech (repetition of the syllable “ba” at a 2 Hz repetition rate) (Keshavarzi et al., 2024a). When administered to children with DD, the rhythmic speech paradigm reliably reveals an atypical preferred phase in the delta band (Power et al., 2013; Keshavarzi et al., 2022b). This could be suggestive of the DD brain encoding a less-accurate representation of low frequency envelope information. Accordingly, by TS theory a similar delta-band impairment could be anticipated in DLD. However, the participants with DLD in Keshavarzi et al. (2024a) who listened to rhythmic speech did not show atypical preferred phase in the delta band. Instead, they showed enhanced theta and low gamma band power during the entrainment period, with significantly greater power than the TD control children. Further, when group differences (DLD vs TD) in resultant phase were computed in exploratory analyses, theta-low gamma resultant phase differed significantly between groups, as did delta-low gamma phase, suggestive of atypical low frequency-high frequency oscillatory dynamics. These data suggest that group differences (DLD vs TD) in oscillatory encoding of rhythmic speech may occur both in the power domain and also when low-frequency phase modulates gamma-band amplitude (PAC) and/or shows altered phase synchrony with gamma-band phase (PPC). Such data are suggestive of a different underlying relationship between low-frequency phase and high-frequency phase during speech listening in the DLD brain, at least when rhythmic speech is the input signal.

In the current EEG study, we measured cortical tracking of low-frequency speech information during a story listening task utilising natural speech, computing backward multivariate temporal response functions (mTRFs, see Crosse et al., 2016) to assess cortical tracking globally (whole brain). We also conducted analyses focused on the right temporal region, as prior DD studies have identified atypical right-lateralised cortical tracking in the delta band in children with dyslexia (Di Liberto et al., 2018). On the basis of TS theory, we predicted impaired decoding of envelope information <10 Hz (delta and theta bands) in the children with DLD, H1. H1 thus tested the primary prediction of TS theory, that children with DLD would exhibit less accurate low-frequency cortical tracking of natural continuous speech, either globally or in a pre-defined right temporal region. As a secondary objective, we also assessed both band-power and cross-frequency coupling relationships between bands. These secondary objectives were motivated by both the prior infant data and the prior TS-driven studies described above (Araújo et al., 2024; Keshavarzi et al., 2024a). As the available resting state oscillatory literature with infants at-risk for DLD and children with DLD identifies theta and gamma resting state differences (Benasich et al., 2008; Choudhury & Benasich, 2011; Gou et al., 2011; Cantiani et al., 2016, 2019; Meng et al., 2022; Lum et al., 2022), we predicted greater theta and gamma power in DLD (H2). On the basis of the infant studies (Attaheri et al., 2024) and the DLD classifier study by Araújo et al. (2024), we also predicted atypical delta-theta and theta-gamma PAC (H3). Finally, given the resultant phase data reported by Keshavarzi et al. (2024a) for rhythmic speech, we also tentatively predicted differences in delta-gamma and theta-gamma PPC for the children with DLD (H4).

## 2. Methods and Material

### 2.1. Participants

Sixteen typically developing children (TD control group, mean age of 9.1 ± 1.1 years) and sixteen children with DLD (mean age of 9 ± 0.9 years) took part in the study. All participants were taking part in an ongoing study of auditory processing in DLD (Keshavarzi et al., 2024a), and those children with suspected language difficulties were nominated by the special educational needs teachers in their schools. Children in the TD group were nominated by classroom teachers as being typically-developing. All children had English as the main language spoken at home. All participants exhibited normal hearing when tested with an audiometer. In a short hearing test across the frequency range 0.25–8 kHz (0.25, 0.5, 1, 2, 4, 8 kHz), all children were sensitive to sounds within the 20 dB HL range. Language status was ascertained by a trained speech and language therapist (the second author), who administered two subtests of CELF-V (Wiig et al., 2017) to all the participants: recalling sentences and formulating sentences. Those children who appeared to have language difficulties then received (depending on age) two further CELF subtests drawn from word structure, sentence comprehension, word classes and semantic relationships. Children who scored at least 1 S.D. below the mean (7 or less when the mean score is 10) on at least 2 of these 4 subtests were included in the DLD group. Oral language skills in the control children were thus measured using only two CELF tasks, recalling sentences and formulating sentences, and all TD control children scored in the normal range (achieving scores of 8 or above). Due to the Pandemic, although all 16 control children received the recalling sentences task, only 11 also received the formulating sentences task (all children with DLD received 4 CELF tasks). The Picture Completion subtest of the Wechsler Intelligence Scale for Children (WISC IV, Wechsler, 2016) was used to assess non-verbal intelligence (NVIQ) and did not differ significantly between groups. Group performance for the tasks administered to both groups is shown in Table 1. All participants and their parents provided informed consent to participate in the EEG study, in accordance with the Declaration of Helsinki. The study was reviewed by the University of Cambridge, Psychology Research Ethics Committee and received a favourable opinion.

**Table 1.** Details of the participating children, showing Scaled Scores (ScS, population mean = 10) and Standard Scores (SS, population mean = 100).

|  | DLD | TD Controls | Significance |
| --- | --- | --- | --- |
| N | 16 | 16 | — |
| Age (months) | 108.3 (11.3) | 108.9 (13.3) | 0.585 |
| Recalling Sentences ScS | 5.3 (1.7) | 10.9 (2.3) | 0.0001 |
| Formulating Sentences ScS | 4.4 (1.9) | 10.8 (1.7) | 0.0001 |
| WISC Picture Completion ScS | 7.9 (3.0) | 9.9 (2.8) | 0.075 |

Note. WISC: Wechsler Intelligence Scale for Children.

### 2.2. Experimental set-up and stimuli

The experimental setup and stimuli utilised in the EEG study were identical to those used in a prior TS-driven study of children with DD, please see Keshavarzi et al. (2022a). The participants listened to a 10-minute story for children, *The Iron Man: A Children’s Story in Five Nights* by Ted Hughes. During the experiment, participants were instructed to listen to the speech carefully and to look at a red cross (+) shown on the computer screen. The sampling rate was 44.1 kHz for presenting the auditory stimulus, and EEG data were concurrently collected at a sampling rate of 1 kHz using a 128-channel EEG system (HydroCel Geodesic Sensor Net). To ensure the children attended to the task, they were informed beforehand that they would be asked two simple comprehension questions after the story. We assessed the correct scores on the two comprehension questions administered after the story-listening paradigm. Comprehension was assessed by scoring these two questions, with possible scores ranging from 0 to 2. Mean scores (± SD) were 1.9 ± 0.3 and 1.81 ± 0.4 for TD and DLD groups, respectively. A Wilcoxon rank-sum test revealed no significant difference between the groups (*z* = 0.45, *p* = 0.65), indicating that comprehension performance was comparable.

### 2.3. Auditory stimuli and EEG data pre-processing

We initially obtained the broad-band envelope by computing the absolute value of the analytical signal from the speech stimuli. The resulting envelope was then band-pass filtered within the frequency range of 0.5–8 Hz, after which the filtered speech envelope was downsampled to 100 Hz. This low-frequency envelope was used for all analyses conducted in this study.

The EEG recordings were first referenced to the Cz electrode. The data were then band-pass filtered between 0.2 and 42 Hz using a zero-phase finite impulse response (FIR) filter as implemented in the MNE-Python software package (Gramfort et al., 2013). EEG signals were subsequently downsampled to 100 Hz for all downstream analyses. Bad channels were identified manually for each participant based on visual inspection of both time-domain waveforms and power spectral characteristics, and were then interpolated using spherical spline interpolation implemented in MNE-Python. Independent Component Analysis (ICA) was subsequently performed using the Infomax algorithm. The resulting independent components (ICs) were manually evaluated and classified as neural or artefactual (e.g., eye blinks, eye movements, and cardiac activity) based on inspection of their scalp topographies, power spectra, and time-domain waveform of the components. Identified artefactual components were removed, and the ICA solution was then applied to the broadband EEG data. For PPC and PAC analyses, band-pass filtering was applied using the MNE-Python library to extract delta (0.5–4 Hz), theta (4–8 Hz), and low gamma (25–40 Hz) frequency bands. For decoding-related computations, MATLAB was used to implement filtering.

### 2.4. Preprocessing quality control

The number of interpolated bad channels did not differ between groups. The control group showed a mean of 9.13 ± 3.16 bad channels per participant, while the DLD group showed a mean of 9.00 ± 2.88 bad channels. A two-tailed independent-samples t-test revealed no significant group difference (*p* = 0.91). Similarly, the number of excluded ICs following ICA artefact rejection was comparable between groups. The control group had a mean of 15.56 ± 2.58 excluded ICs, whereas the DLD group had a mean of 14.88 ± 2.33 excluded ICs. This difference was not statistically significant (*p* = 0.44). These results indicate that data quality and preprocessing procedures were well matched across groups, and that subsequent group differences in neural measures cannot be attributed to systematic differences in channel quality or ICA-based artefact rejection.

### 2.5. Computation of band-power

To investigate the power of neural response over the entrainment period used for analysis, we calculated the power of EEG responses for each child and group through a four-step process: (1) Individual EEG channel power calculation using the *pspectrum*() function in MATLAB, applying Welch’s method (Welch, 1967); (2) Power calculation by averaging across channel power values from step 1; (3) Participant power calculation by averaging across power values from step 2 for each participant; (4) Group power calculation by averaging across power values from step 2 for all children within each group.

### 2.6. Spectral parameterisation

Recent advances in EEG signal processing have revealed that neural power spectra contain both periodic and aperiodic components, both of which may have functional significance for behaviour (Donoghue et al., 2020). To distinguish narrowband oscillatory activity from broader spectral characteristics in our data, we therefore additionally applied spectral parameterisation to separate periodic and aperiodic components of the power spectrum. Analyses were conducted using the python FOOOF package (https://fooof-tools.github.io/).

Spectral parameterisation was performed using the FOOOF algorithm (Donoghue et al., 2020) applied to the channel-averaged power spectral density (PSD) for each participant, computed using Welch’s method. FOOOF models the spectrum as the sum of an aperiodic component (parameterised by offset and exponent) and Gaussian peaks reflecting periodic oscillatory activity. The aperiodic component was modelled using a fixed (no-knee) model.

After fitting, we extracted the aperiodic parameters (offset and exponent) and derived the aperiodic-adjusted (“flattened”) spectra by removing the fitted aperiodic component. Band-limited activity (delta, theta, and low gamma) was then recomputed from these flattened spectra to isolate periodic contributions independent of broadband spectral slope. Group comparisons were repeated on these aperiodic-adjusted measures to determine whether observed effects in conventional band-power analyses reflected oscillatory activity or broader spectral characteristics.

### 2.7. Computation of cross frequency PAC

Cross frequency PAC refers to the modulation of the amplitude of the signal in a high frequency band by the phase of a low frequency oscillation. Here we quantified the strength of this type of modulation using the modulation index (*MI*; Tort et al. 2008):

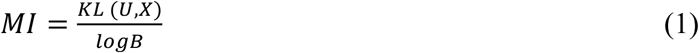

where *B* (=18) is the number of bins, *U* refers to the uniform distribution, *X* is the distribution of the data, and *KL* (*U*, *X*) is Kullback–Leibler distance which is calculated as:

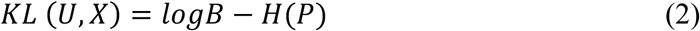

where *H*(.) is the Shannon entropy and *P* is the vector of normalised averaged amplitudes per phase bin which is calculated as:

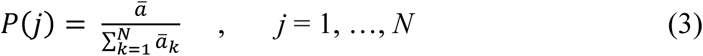

where ā is the average amplitude of each bin, *k* refers to running index for the bins. Note that *P* is a vector with *N* elements.

### 2.8. Computation of cross frequency PPC

Cross-frequency PPC refers to the phase synchrony between oscillations in two frequency bands. To quantify the PPC, we employed the phase-locking value (*PLV*; Lachaux et al., 1999), which is calculated as:

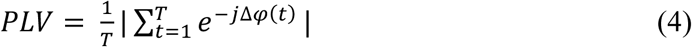

where |.| refers to the absolute value operator, *T* is the number of time samples, and Δφ(*t*) is the phase difference, and is calculated as:

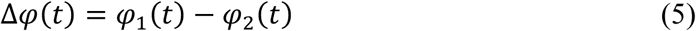

where φ_1_(*t*) is the instantaneous phase of first signal and φ_2_(*t*) is the instantaneous phase of the second signal. Although both PAC and PPC assess cross-frequency interactions, PAC quantifies phase-amplitude modulation whereas PPC quantifies phase synchrony between oscillations. Accordingly, different metrics were used to appropriately capture these distinct coupling mechanisms.

### 2.9. Stimulus reconstruction using backward TRF model

We applied the backward TRF model (Crosse et al., 2016) to reconstruct the stimuli envelopes from the EEG signals at each frequency band of interest (delta, theta, alpha), and to estimate decoding accuracy. This was done for both the whole head (global comparison) and for a pre-determined region of the right hemisphere. The right-hemisphere temporal region of interest (ROI) was defined a priori based on electrode locations overlying the right superior temporal area in the EGI HydroCel 128-channel layout. The following electrodes were included in this ROI: E100, E102, E109, E114, E121, E116, E101, E107, E108, E115, E113, and E120.

The backward TRF model predicts (reconstructs) the stimulus envelope, denoted as *s*(*t*), through the following equation:

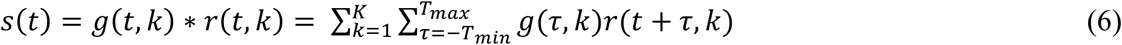

where *g*(τ, *k*) is a decoder representing the linear mapping from the neural response to the stimulus envelope for time lag τ and channel *k*. Parameters *T_min_* = 0ms and *T_max_* = 400 *ms* determine the minimum and maximum time lags, respectively, and *r*(*t*, *k*) denotes the neural response at channel *k*. As an additional exploratory analysis, reconstruction accuracy was computed for multiple maximum lag values (*T_max_ =* 0-500ms) to characterise how decoding performance varied with the temporal integration window of the backward TRF model. In this framework, *T_max_* defines the cumulative interval over which neural activity *T_min_* = 0ms to *T_max_* contributes to stimulus reconstruction. These analyses were exploratory. The validation procedure utilised to regularise the backward linear model was the “leave-one-out” cross-validation (using *mTRFcrossval* function from the mTRF Toobox) in which one trial is “left out” or used for testing and the remainder are used to train the model. Each trial was 1 minute in length. This process is repeated across all trials. The performance of the model was evaluated using the Pearson correlation between the estimated envelope and the actual envelope, with a higher correlation value indicating more accurate decoding by the model.

### 2.10. Between-group backward TRF analysis

To simulate normative decoding of the speech envelope from the neural responses at each frequency band, we used EEG recorded from the TD control children. For each frequency band of interest, we used a randomly selected subset of control children (*C_train_* = 8) for training 8 backward TRF models. A final model was obtained by averaging across all 8 individual trained models. Subsequently, this averaged model was independently tested on the remaining TD control children (*C_test_* = 8) as well as on 8 randomly selected children with DLD. We then computed the mean correlation values for both the tested TD control children and the children with DLD, resulting in a single correlation value for each group. This procedure was iterated 100 times, employing random permutations across all control children, thereby enabling the use of different combinations of 8 TD control children (for *C_train_* and *C_test_*) for training and testing the model. As a result, 100 normative (averaged) models were generated. Accordingly, 100 correlation values were produced for both the TD control children and the children with DLD, yielding an averaged correlation value for each group. This approach facilitated the assessment of potential differences in speech decoding accuracy at the group level between children with and without DLD. Thus, the between-group analysis used a group-derived normative TRF model.

The rationale behind the between-group backward TRF analysis is to establish a normative decoding model based on TD children, which serves as a reference for assessing neural decoding performance in children with DLD. By averaging TRF models trained on random subsets of TD children, we create a generalised normative model that captures typical neural encoding of speech. Testing this model on DLD children allows us to assess how well their neural responses align with normative patterns, providing insights into potential deviations in speech-brain coupling. This generalisation of the normative model to DLD data is a key feature of the analysis, as it enables us to evaluate the extent to which DLD children’s neural responses conform to or diverge from typical encoding patterns.

### 2.11. Within-child backward TRF analysis

In order to examine the consistency of perceptual experience regarding low-frequency envelope information within the speech signal for individual children, we constructed a single backward TRF model for each child in the study, utilising only the data from that specific child. This within-child approach allowed us to investigate whether the representations of speech developed by individual children with DLD exhibit lower consistency than those developed by individual TD children. To accomplish this, we allocated 80% of the data from each child for training the TRF model, reserving the remaining 20% for testing the model. Subsequently, we computed the Pearson correlation between the estimated speech envelope and the actual speech envelope for each child individually and for each frequency band. This yielded a single correlation score for each band for every child. These scores were then aggregated by group (TD, DLD) for each frequency band and compared between groups. Thus, the within-child analysis used fully subject-specific TRF models. This approach allowed us to assess whether there are differences in the consistency of speech-based representations between children with and without DLD. This within-child analysis was conducted independently for the whole brain and for the right temporal region.

### 2.12. Computation of chance level for the between-group backward TRF model analysis

To assess the statistical significance of stimulus reconstruction accuracy as estimated by the between-group backward TRF model analysis, null models were computed for each frequency band of interest. To generate these null models, eight TD control children were randomly selected for training 8 backward TRF models, and EEG data were permuted across different story sections for each of these TD control children. A final model was obtained by averaging across all 8 individual trained models. Subsequently, this averaged model was independently tested on the remaining TD control children (*C_test_* = 8). We then computed the mean correlation values for the tested children, resulting in a single correlation. This process was repeated 200 times for each frequency band to generate probability density functions, which serve as statistical measures determining the probability distribution of a variable.

### 2.13. Computation of chance level for the within-child backward TRF model analysis

In order to assess the statistical significance of stimulus reconstruction accuracy as estimated by the within-child backward TRF model analysis, null models were computed for each frequency band of interest using data from TD control children. To generate these null models, all sixteen TD control children were considered. For each permutation iteration, a backward TRF model was constructed separately for each TD child. In each iteration, 80% of the data were used for training and 20% for testing, as in the main analysis. However, for the null model, the correspondence between EEG data and speech envelope was disrupted by permuting the EEG data across different story sections within the training set, thereby removing the true stimulus-response relationship while preserving the overall spectral characteristics of the data. The Pearson correlation between the estimated speech envelope and the actual speech envelope was then calculated separately for each TD child and each frequency band within these null models, resulting in a single correlation value for each band and child. For each permutation iteration, these child-level correlation values were averaged across the sixteen TD children to yield one mean reconstruction accuracy value per frequency band. This entire procedure was repeated 200 times per frequency band, and the 200 mean reconstruction accuracy values were used to construct the null distribution (probability density function). The within-child chance level was conducted independently for the whole brain and for the right temporal region.

### 2.14. Statistical analysis

Given the relatively small sample size in each group (*n* = 16), and because several neural measures exhibited non-normal distributions when assessed using Shapiro-Wilk tests, we employed non-parametric statistical analyses for most of our group comparisons. Specifically, Wilcoxon rank-sum tests (two-tailed) were used to compare TD and DLD groups for band-power, PAC, and PPC measures.

For backward TRF decoding analyses, two types of comparisons were conducted. First, decoding accuracy for each frequency band was compared against chance by evaluating performance relative to null models. Second, group comparisons of decoding accuracy used either Wilcoxon rank-sum tests or two-sample t-tests when the distribution of correlation values (e.g., across the 100 backward TRF models) met normality criteria. This approach ensured continuity with prior mTRF research while retaining statistical robustness. In addition, Bayesian factor analysis was employed for within-child decoding comparisons to quantify evidence for or against group differences, providing a more informative interpretation of null results.

## 3. Results

### 3.1. Does the accuracy of speech envelope decoding vary between children with and without DLD?

To investigate the first prediction H1 concerning the expected impaired accuracy of low-frequency speech envelope decoding in children with DLD, the between-group backward TRF analysis method (as detailed in Section 2.10) was independently applied for each frequency band. To assess the statistical significance of the stimulus reconstruction accuracy achieved for each frequency band by group, we first compared decoding accuracy in each band to that of the null models (which represent the chance level for each band, as described in Section 2.12). The between-group analysis revealed that stimulus reconstruction accuracy was significantly higher than chance level for both groups of children in all three bands (Figure S1). To compare the decoding accuracy between the TD and DLD groups, we conducted independent two-sample t-tests. The results indicated that reconstruction accuracy did not differ between the TD and DLD groups in any frequency band (Figure 1; delta: *p* = 0.22; theta: *p* = 0.25; alpha: *p* = 0.12). Accordingly, global (whole brain) cortical tracking of continuous speech estimates did not differ between groups, contrary to H1. However, the modest sample size may have limited our statistical power to detect subtle or spatially distributed group differences in cortical tracking.

**Figure 1.**
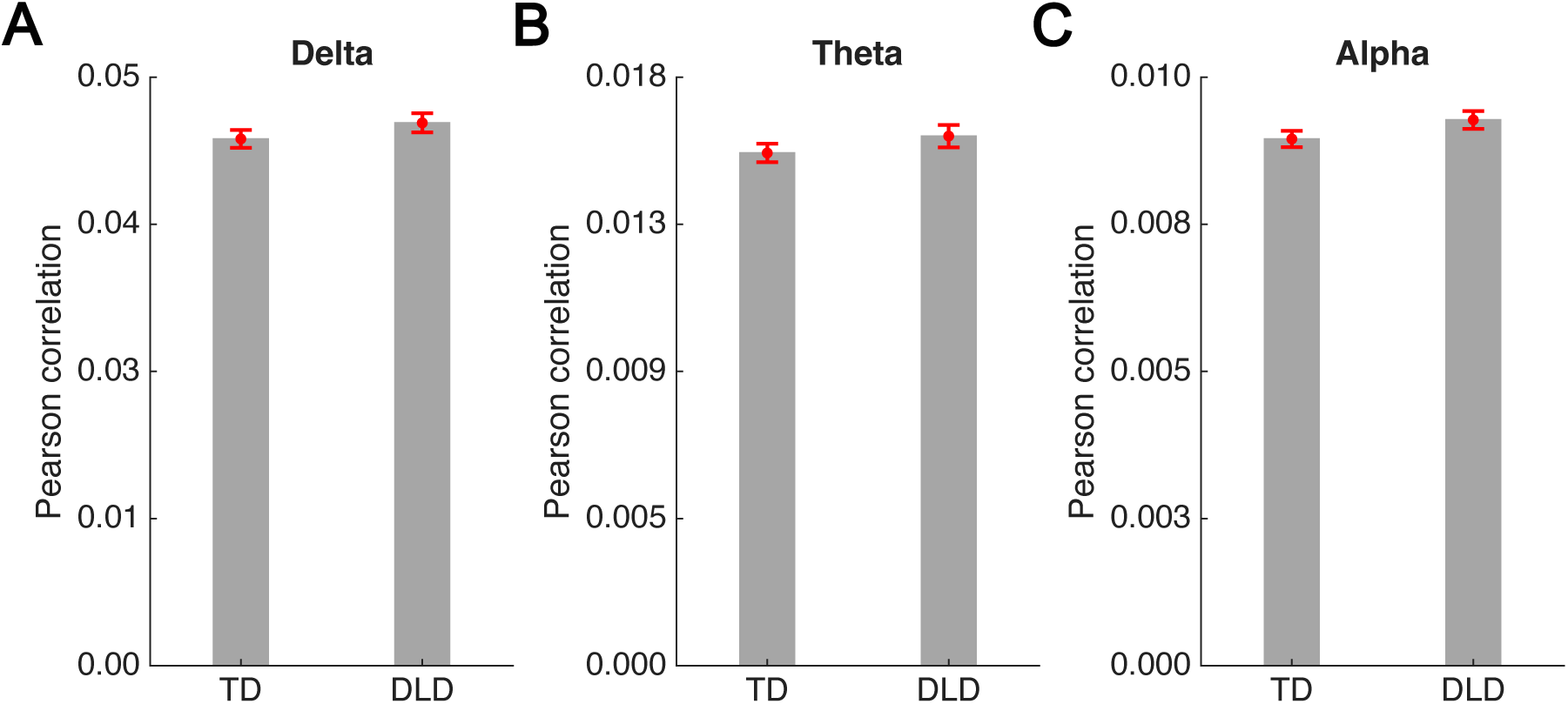
Stimulus-reconstruction accuracy in for the delta band (Panel A), theta band (Panel B), and alpha band (Panel C) for children in the DLD and TD groups, based on the between-group analysis. Bars represent mean Pearson correlation averaged across 100 backward TRF models. Error bars indicate the standard error of the mean.

To examine whether decoding performance varied with the size of the temporal integration window, we ran an exploratory analysis using different values of the maximum time lag parameter (*T_max_* = 0 ms, 100 ms, 200 ms, 300 ms, 400 ms, and 500 ms). Figure S2 presents the decoding accuracy values obtained at each value of *T_max_* No group differences were found at any individual *T_max_* value.

### 3.2. Do individual children with DLD develop less precise representations of low-frequency information in speech?

To investigate whether individual children with DLD exhibit less precise representations of the low-frequency envelope information in the speech signal compared to individual TD children, the within-child backward TRF analysis method (as described in Section 2.11) was applied independently for each frequency band and compared by group. Comparisons with the null models revealed that stimulus reconstruction accuracy surpassed chance levels for both groups across all three frequency bands (Figure S3). Figure 2 displays the box plots illustrating stimulus reconstruction accuracies for both the TD and DLD groups, categorised by band, based on the backward TRF models from individual children. To compare the reconstruction accuracy between the two groups, we conducted independent two-sample t-tests. The results revealed no significant difference between the TD and DLD groups across any band (delta band, Figure 2A, *p* = 0.95; theta band, Figure 2B, *p* = 0.87; alpha band, Figure 2C, *p* = 0.83). Additionally, Bayesian factor analysis was employed to quantify the strength of evidence for the alternative model. The findings indicated greater evidence in favour of the null model for all three bands (Bayesian factor analysis; *BF10s* > 0.26). Accordingly, within-child cortical tracking of continuous speech at the whole-brain level did not differ between groups, contrary to H1. As an exploratory analysis, we additionally calculated within-child decoding accuracy across multiple *T_max_* values (0, 100, 200, 300, 400, and 500 ms) using whole-brain data. Figure S4 presents reconstruction accuracies for the TD and DLD groups at each temporal window separately for the delta, theta, and alpha bands. No group differences were found at any individual *T_max_* value.

**Figure 2.**
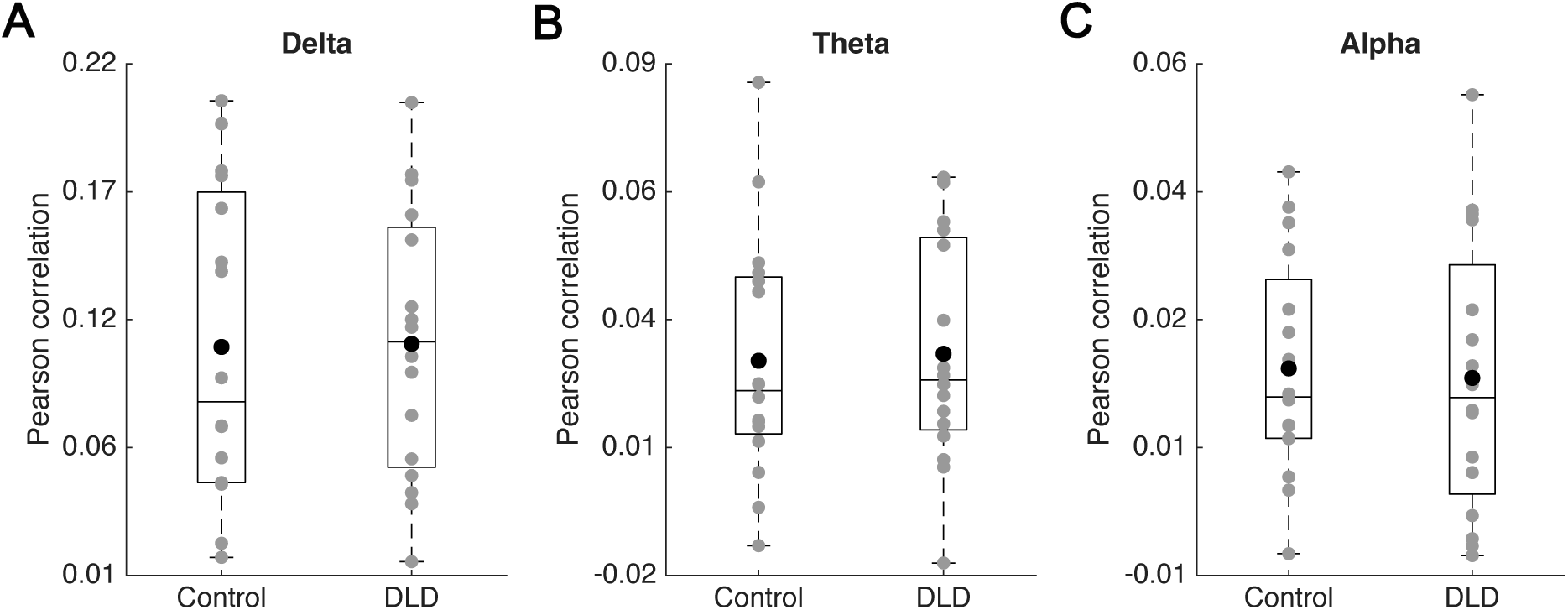
Stimulus-reconstruction accuracy for children in the DLD and TD groups, in the delta (Panel A), theta (Panel B), and alpha (Panel C) bands, based on the within-child analysis. Grey circles represent the reconstruction accuracy scores for individual children, while the black circles indicate the mean accuracy scores for each band and group.

However, prior TS-driven studies suggest that the right hemisphere may be important regarding impaired cortical tracking in DD (Di Liberto et al., 2018). Accordingly, we also compared decoding accuracy between the two groups within the right temporal region. The within-child backward TRF analysis was again conducted separately for each frequency band. For both TD and DLD children, stimulus reconstruction accuracy exceeded chance levels across all three frequency bands (Figure S5). Figure 3 presents box plots of reconstruction accuracy for the TD and DLD groups, shown separately for each band and based on individual children’s backward TRF models. Group differences in reconstruction accuracy were assessed using independent two-sample t-tests. The results revealed a significant difference between the TD and DLD groups in the delta band (*p* = 0.03), but no significant differences were observed in the theta (*P* = 0.46) or alpha (*p* = 0.70) bands. Accordingly, H1 was supported regarding the right temporal region.

**Figure 3.**
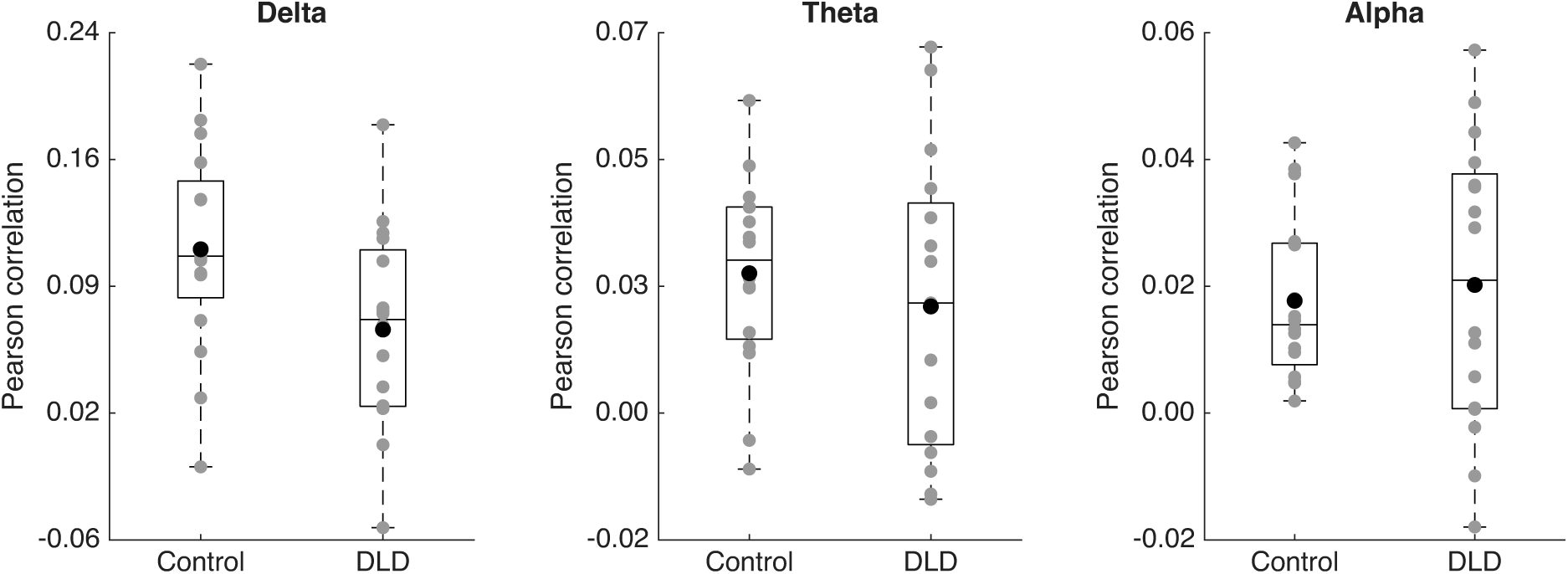
Stimulus-reconstruction accuracy within the right temporal region for children in the DLD and TD groups, in the delta (Panel A), theta (Panel B), and alpha (Panel C) bands, based on the within-child analysis. Grey circles represent the reconstruction accuracy scores for individual children, while the black circles indicate the mean accuracy scores for each band and group.

As an additional exploratory analysis, decoding accuracy was calculated for different temporal integration windows by investigating different values of *T*_;->_. Figure S6 presents the reconstruction accuracy values for multiple *T_max_* values (0, 100, 200, 300, 400, and 500 ms) for each frequency band for each group separately. In the delta band, maximum lag values *T_max_* of 200, 300, 400, and 500 ms all showed significantly higher reconstruction accuracy in the TD group relative to the DLD group (uncorrected).

### 3.3. Comparison of band-power between groups

To investigate H2, we compared the band-power between groups for the delta, theta and gamma bands. Wilcoxon rank sum tests were applied for each frequency band after removing outliers (3 outliers in TD and 3 outliers in DLD). A data point was classified as an outlier if its corresponding power value exceeded 1.5 times the interquartile range above the upper quartile or fell below the lower quartile of the group’s population data. Figure 4 shows the average power versus frequency separately for each group and for delta (see Figure 4A), theta (see Figure 4B), and low gamma (see Figure 4C) bands. A significant difference between TD control children and children with DLD was found in the theta (*z* = –2.26, *p* = 0.02) and low gamma (*z* = –2.92, *p* = 0.003) bands but not in the delta band (*z* = –1.64, *p* = 0.10).

**Figure 4.**
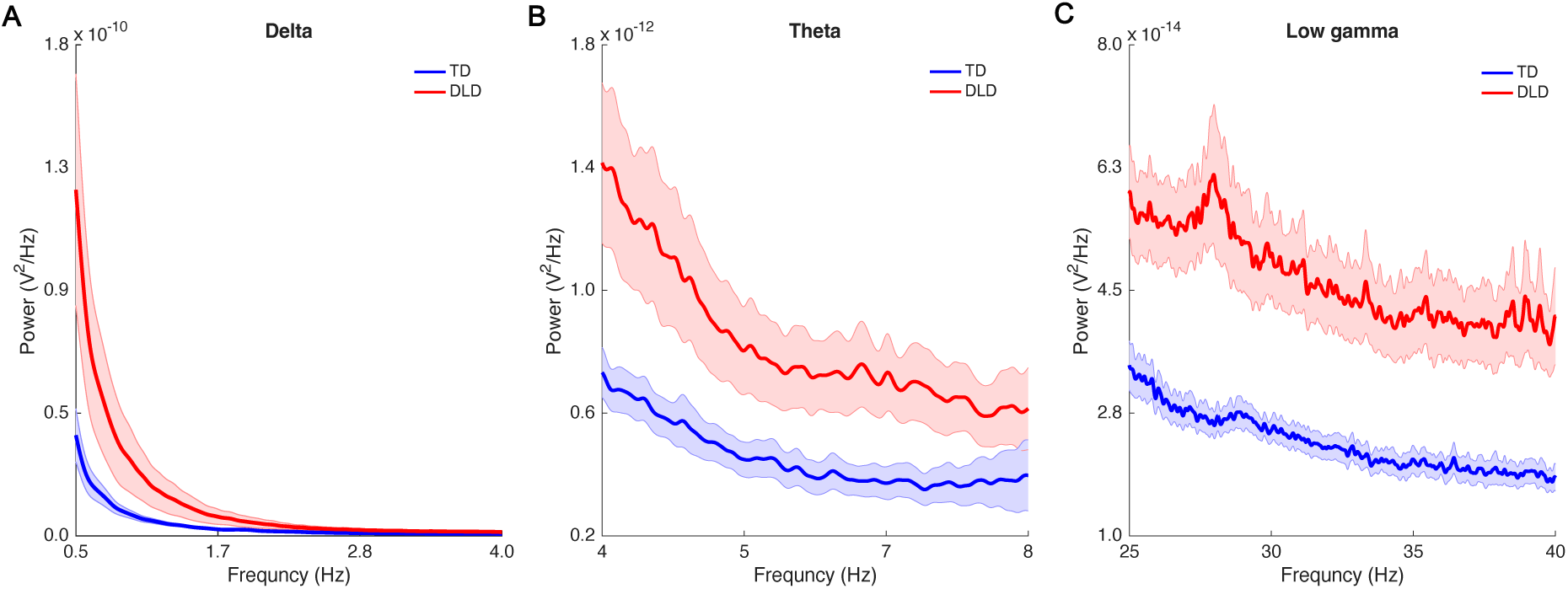
Mean power for delta-band (Panel A), theta-band (Panel B), and low gamma-band (Panel C). The blue and red curves represent the power density function for the TD and DLD groups, respectively. The shaded areas show the standard error of the mean for each group.

To determine whether these group differences reflected narrowband oscillatory activity or broader spectral characteristics, we additionally applied spectral parameterisation to separate periodic and aperiodic components of the power spectrum (see Section 2.6, Donoghue et al., 2020). Following removal of the aperiodic component, group comparisons were repeated on the aperiodic-adjusted (flattened) spectra using the same non-parametric statistical approach. Under this control analysis, no significant group differences were observed in aperiodic-adjusted delta, theta, or low gamma activity (all *p* > 0.37). This indicates that the group differences observed in conventional band-power measures are primarily driven by broadband spectral components and not by oscillatory power per se. Accordingly, these oscillatory power differences are not analysed further.

### 3.4. Comparison of PAC between groups

To test H3, that delta-theta and theta-gamma PAC may be atypical in children with DLD, we compared the PAC measures between the groups. Wilcoxon rank sum tests were applied for each PAC measure after removing outliers. The results revealed that there was no significant difference between the two groups in delta-theta PAC (*z* = –0.11, *p* = 0.91; 1 outlier in TD and 2 outliers in DLD), nor in delta-low gamma PAC (*z* = 0, *p* = 1; 1 outlier in TD and 1 outlier in DLD), nor in theta-low gamma PAC (*z* = –1.56, *p* = 0.12; 1 outlier in DLD). These PAC values are depicted as bar plots in Figure S7.

### 3.5. Comparison of PPC between groups

To test H4, that PPC may differ between groups, we applied two-sample t-tests to compare PPC after removing outliers. The results showed no significant difference between the two groups neither for delta-theta PPC (*p* = 0.72), nor for delta-low gamma PPC (*p* = 0.21; 1 outlier in TD and 1 outlier in DLD), nor for theta-low gamma PPC (*p* = 0.09). These PPC values are depicted as bar plots in Figure S8.

## 4. Discussion

The current study explored neural processes during continuous speech listening in two groups of children aged on average 9 years, who were classified as either TD or DLD. Given the extensive nature of the language difficulties exhibited by children with DLD (CATALISE, Bishop et al., 2017), we combined TS theory with a broader oscillatory framework, and explored both low-frequency cortical tracking of the speech signal by oscillatory responses in the delta and theta bands, and band-power, PAC and PPC between oscillatory networks responding at different temporal rates (Giraud and Poeppel, 2012; Gross et al., 2013). The EEG data collected here in a continuous speech listening paradigm provided partial support for the impairments in low-frequency cortical tracking predicted by TS theory. The key prediction (H1) that delta and/or theta band cortical tracking of the speech signal should be less accurate in children with DLD was supported in the right hemisphere for the delta band. In the within-child mTRF analyses, the children with DLD showed equivalent reconstruction accuracy to the TD children for all frequency bands measured (delta, theta, alpha) globally (whole brain), but showed significantly reduced accuracy for the delta band in the right temporal region. In extra exploratory analyses confined to this region, significant group differences were observed for *T_max_* values of 200, 300, 400, and 500 ms (uncorrected) in the delta band. As these integration windows are cumulative, they do not represent independent latency-specific effects, rather they are supportive of the main predefined decoding result for the 0–400 ms window. Accordingly, at least using the mTRF measure of low-frequency cortical tracking employed here, the brains of children with DLD do seem to exhibit altered temporal integration of low-frequency speech information in the right hemisphere.

Regarding the hypothesis about group differences in band power in the theta and gamma bands (H2), both theta power and gamma power were found to be significantly elevated in the DLD group during continuous speech listening prior to application of spectral parameterisation (FOOOF/specparam). This appeared supportive of H2. However, once the periodic (oscillatory) and aperiodic (broadband) components of the power spectrum were separated by spectral parameterisation, the group differences in power were no longer significant. This suggests that the narrowband oscillatory activity typically associated with cognitive processes such as attention, memory, and perception did not differ between the groups studied here. As childhood aperiodic activity in the EEG signal has only been studied in autism spectrum disorder and attention deficit hyperactivity disorder to date (Petri et al., 2025), with a focus on the alpha band, the significance of the current finding remains to be determined. Regarding aperiodic components, Donoghue et al. (2020) showed that aperiodic parameters can have functional significance using a working memory task. In their study, an aperiodic-adjusted alpha power model performed better as a predictor of working memory performance compared to a model using canonical alpha measures, for older but not for younger adults. As the aperiodic-adjusted power models tested here did not in fact differentiate our groups, power differences were not analysed further. However, it is possible that the previous infant and child resting state data regarding DLD (in which both theta and gamma resting state power are elevated in either TD children with poorer language skills or in infants at family risk for DLD) may also change following spectral parameterisation (Benasich et al., 2008; Choudhury and Benasich, 2011; Gou et al., 2011; Cantiani et al., 2016, 2019; Meng et al., 2022, Lum et al., 2022; see also Hill et al., 2022).

Regarding H3 and H4, there was only partial support for the predictions concerning neural dynamics during speech listening that were drawn from a broader oscillatory framework (PAC and PPC). Regarding H3, it had been tentatively predicted that theta-gamma PAC (following Attaheri et al., 2024) and delta-theta PAC (following Araújo et al., 2024) should be atypical in children with DLD during continuous speech listening. Neither prediction was supported, as the average modulation indices did not differ statistically regarding the TD and DLD brains (Figure S7). H3 was also generated in part by data from the Cambridge BabyRhythm infant study, in which a higher theta-gamma modulation index predicted better language outcomes (Attaheri et al., 2024). This finding led here to the prediction that children with DLD may have lower theta-gamma modulation indices. This was not supported, as Figure S7C shows a somewhat higher modulation index for theta-gamma PAC at the group level. Finally, H4 was that PPC may differ between groups, at least when a low-frequency phase has to couple with a high-frequency phase (here tested by delta-gamma and theta-gamma PPC). The PPC analyses (Figure S8) did not support this prediction. The TD control children and the children with DLD showed statistically comparable coupling indices.

The current study has a number of limitations. Firstly, the sample size was small, with 16 children with DLD and 16 control children. This sample size is representative of the existing neural DLD literature (e.g., Heim et al. 2011 studied 17 children with DLD, Nora et al. 2024 studied 17 children with DLD; Choudhury and Benasich 2011 studied 17 infants at risk for DLD). Nevertheless, the modest sample size may have limited statistical power to detect subtle or spatially distributed group differences in cortical tracking, and null findings at the whole-brain level should therefore be interpreted cautiously. In particular, complex neural measures such as mTRF reconstruction, PAC, and PPC may require larger samples to reliably detect small-to-moderate group effects, and replication in larger cohorts will be necessary to establish the robustness and generalisability of the present findings. Secondly, EEG is a relatively noisy neural measure when used with children. More precise data regarding cortical tracking of continuous speech could be gathered by using MEG rather than EEG. Finally, the mTRF is only one method that we used for estimating cortical tracking of the speech signal, and different results may have been found if a different method had been utilised. The mTRF-based reconstruction accuracy estimate relies on linear modelling assumptions and correlation-based metrics, which may not capture more subtle or nonlinear differences. Accordingly, future studies employing converging analytical approaches and larger samples will be important to determine whether the present null findings in whole-brain cortical tracking and cross-frequency coupling reflect genuine neurophysiological equivalence or limited experimental sensitivity. Nevertheless, in prior TS-driven natural language infant studies, low-frequency cortical tracking has been estimated using 3 different measures, the mTRF measure, mutual information, and convolutional neural networks (Keshavarzi et al., 2024b). For TD infants, the pattern of low-frequency cortical tracking using all 3 measures has been broadly similar across all three measures (see Table 1, Keshavarzi et al., 2024b).

In conclusion, the present findings provide partial support for the TS-driven prediction that low-frequency cortical tracking of natural continuous speech should be less accurate in children with DLD. While low-frequency cortical tracking was not globally reduced in our sample at the whole-brain level, children with DLD showed significantly reduced temporal integration of low-frequency speech information in the delta band in the right temporal cortex. As TS theory originally proposed right-lateralised deficits (Goswami, 2011), this finding is consistent with the prediction that atypical low-frequency temporal sampling mechanisms may contribute to language difficulties. It is also interesting to observe that the absence of global impairments in stimulus reconstruction may distinguish the developmental disorders of DLD and DD. Prior TS-driven studies of DD in 3 languages report widespread low-frequency tracking deficits (Molinaro et al., 2016; Destoky et al., 2020; Mandke et al., 2022). It is therefore possible that TS-related mechanisms may operate in DLD in a more spatially constrained and potentially qualitatively distinct manner compared to DD. Future studies, ideally longitudinal and employing larger samples as well as converging methods for estimating cortical tracking, are required to determine how low-frequency temporal sampling and oscillatory dynamics interact in shaping language development in DLD.

## Author contributions

M.K. conceptualisation, methodology, data collection, data analyses, visualisation, writing – original draft; S.R. data collection, writing – review & editing; G.F. data collection, writing – review & editing; L.P. data collection, writing – review & editing; U.G. funding acquisition, project administration, supervision, conceptualisation, methodology, writing – original draft.

## Acknowledgements

The authors would like to thank all the children, families and schools involved in the study. This research was funded by a donation to UG from the Yidan Prize Foundation. The sponsor played no role in the study design, data interpretation nor writing of the report.

## Conflict of Interest

None of the authors have potential conflicts of interest to be disclosed.

---

**Figure S1.**
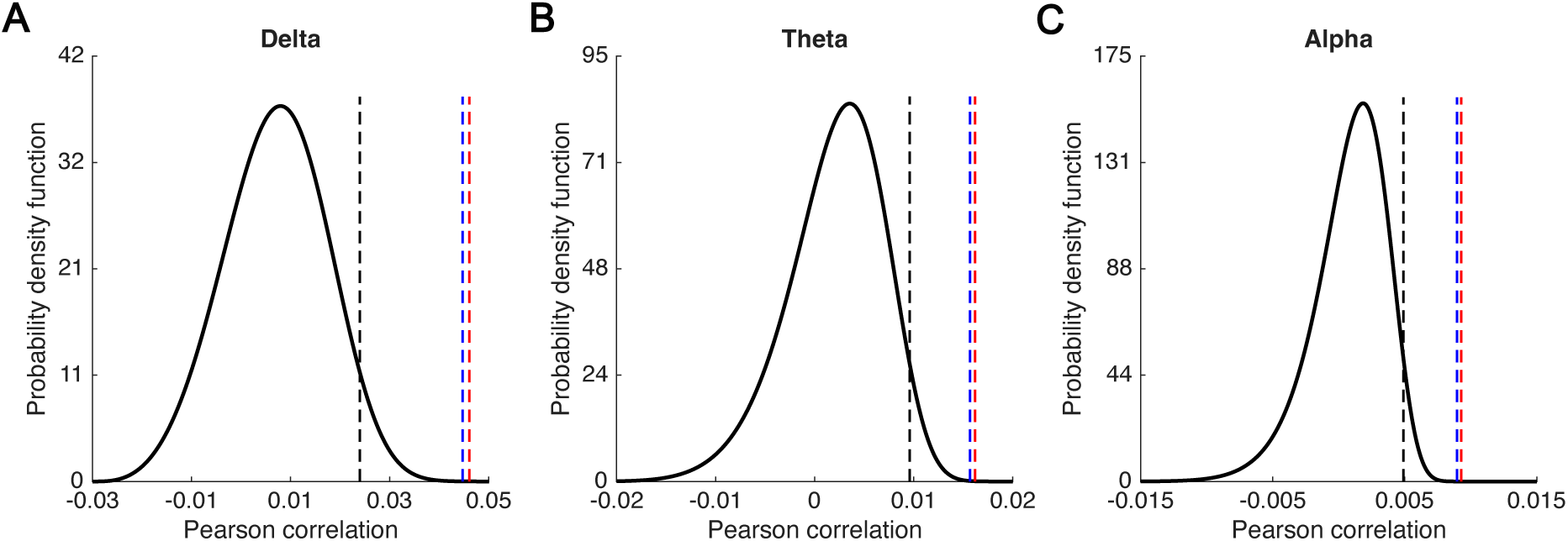
Statistical significance of between-group stimulus reconstruction accuracy for the delta (Panel A), theta (Panel B) and alpha (Panel C) bands summarised by group (Blue, TD group; Red, DLD group). The probability density functions of the null models are represented by the black curves. The dashed black lines indicate the correlation values corresponding to statistical significance (*p* = 0.05). The blue and red dashed lines represent the mean correlation values from the Between-group analysis for the TD and DLD groups, respectively.

**Figure S2.**
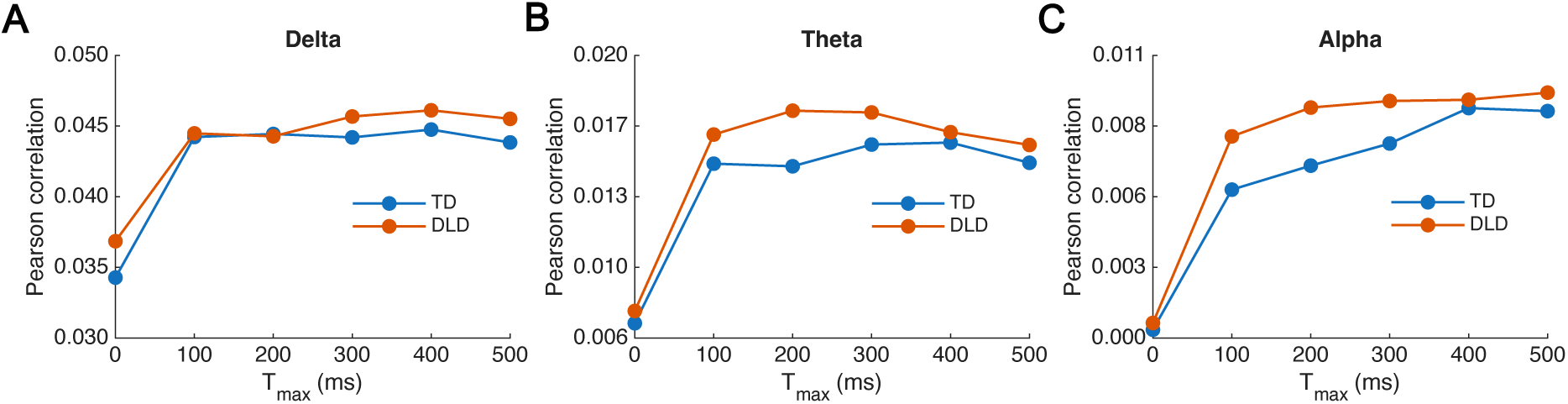
Between-group backward TRF reconstruction accuracy across delays (whole brain). Stimulus reconstruction accuracy as a function of temporal delay (0–500 ms) for the between-group backward TRF model. Panels show (A) delta, (B) theta, and (C) alpha bands. The normative model was trained on TD children and tested separately on TD (blue) and DLD (orange) groups.

**Figure S3.**
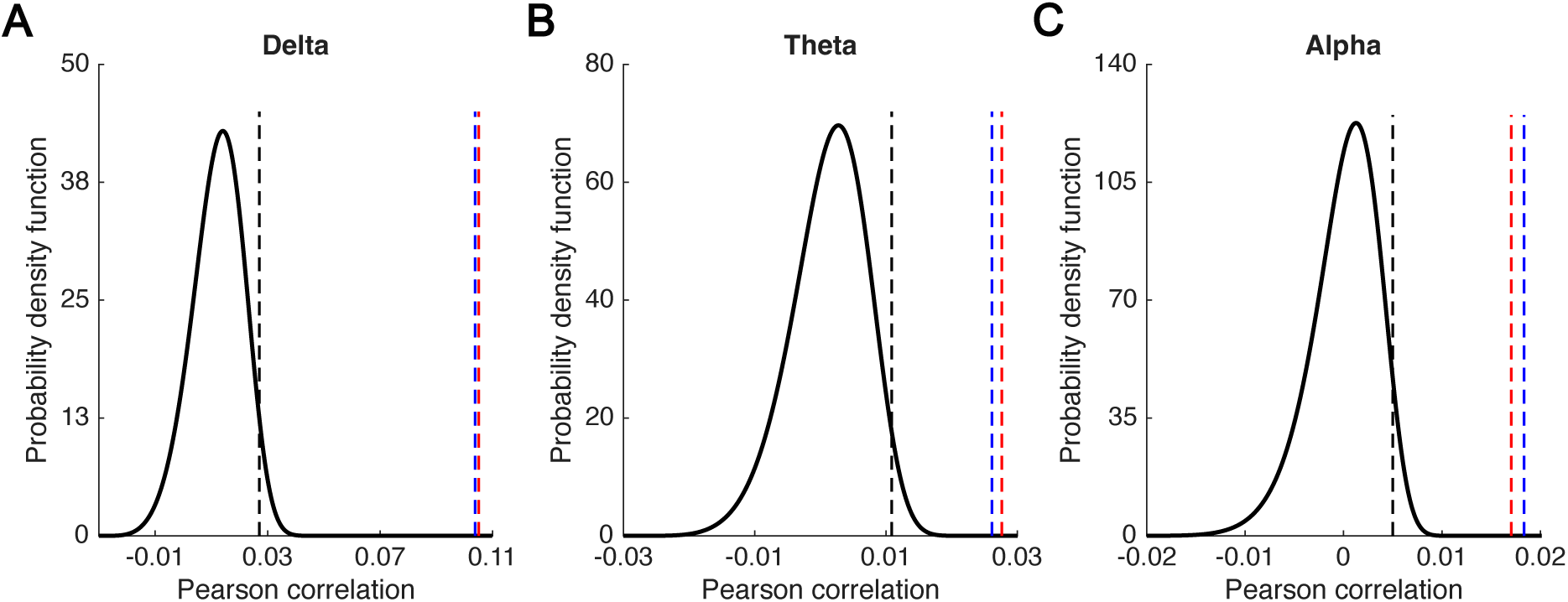
Statistical significance of within-child stimulus reconstruction accuracy (whole brain) for the delta (Panel A), theta (Panel B) and alpha (Panel C) bands summarised by group (Blue, TD group; Red, DLD group). The probability density functions of the null models are represented by the black curves. The dashed black lines indicate the correlation values corresponding to statistical significance (*p* = 0.05). The blue and red dashed lines represent the mean correlation values from the Within-child analysis for the TD and DLD groups, respectively.

**Figure S4.**
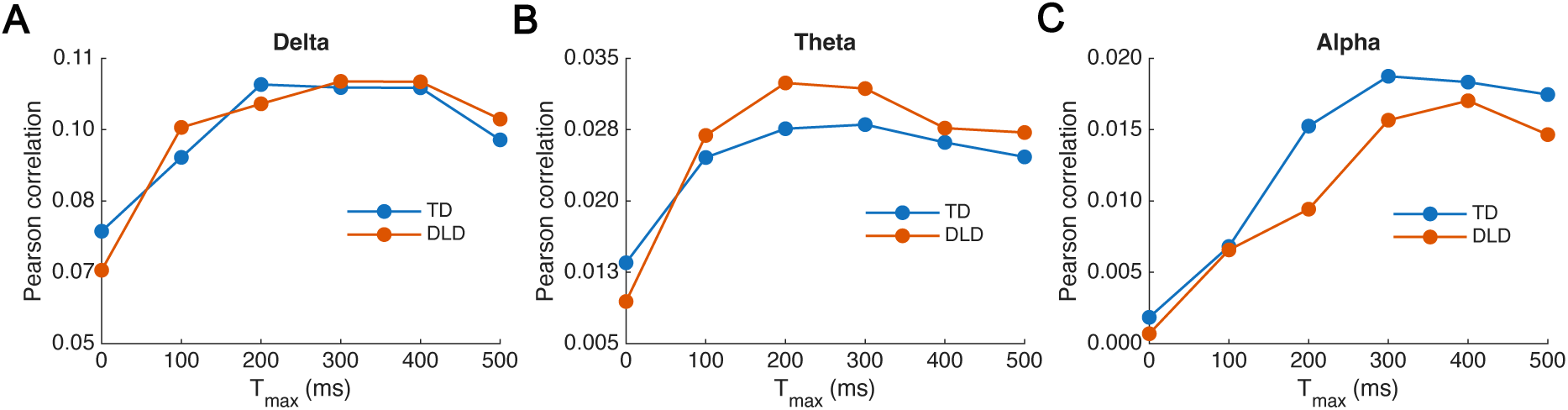
Within-child backward TRF reconstruction accuracy across delays (whole brain). Stimulus reconstruction accuracy as a function of temporal delay (0–500 ms) for within-child backward TRF models computed across the whole head. Panels show (A) delta, (B) theta, and (C) alpha bands. Values represent group-averaged decoding accuracy for TD (blue) and DLD (orange) children.

**Figure S5.**
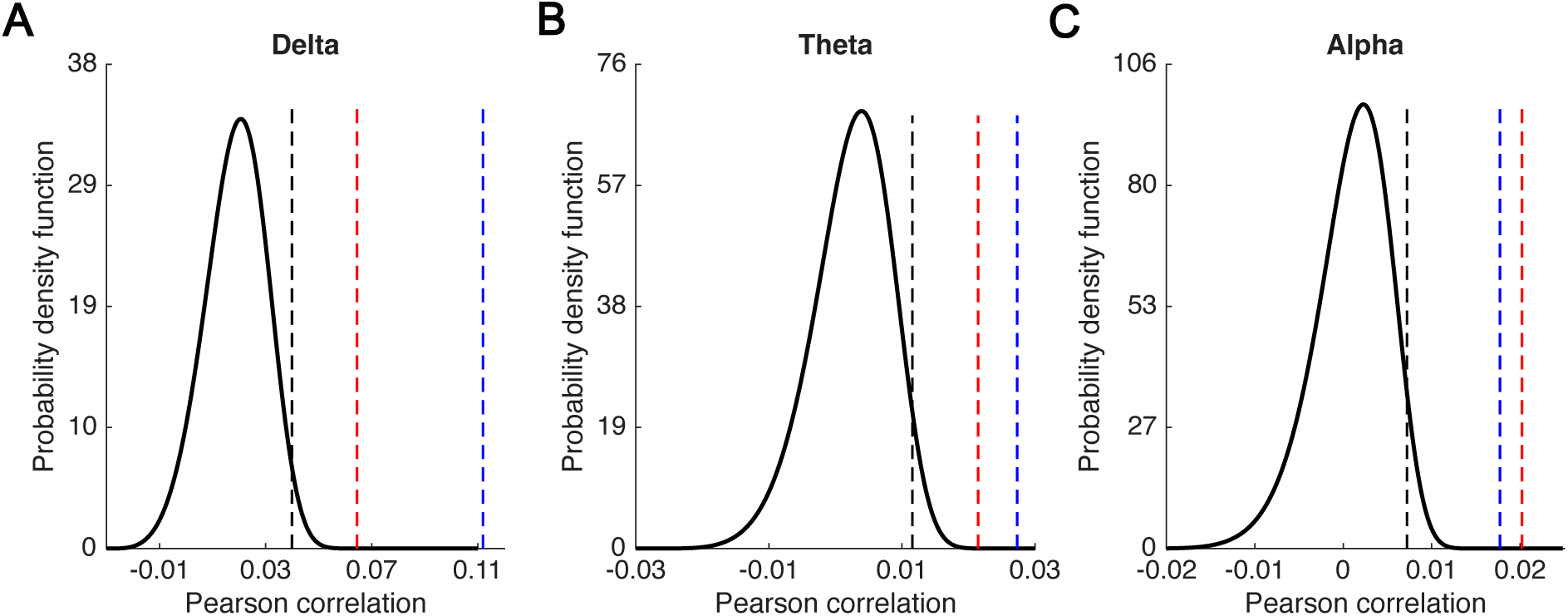
Statistical significance of within-child stimulus reconstruction accuracy for the right temporal region for delta (Panel A), theta (Panel B) and alpha (Panel C) bands summarised by group (Blue, TD group; Red, DLD group). The probability density functions of the null models are represented by the black curves. The dashed black lines indicate the correlation values corresponding to statistical significance (*p* = 0.05). The blue and red dashed lines represent the mean correlation values from the Within-child analysis for the TD and DLD groups, respectively.

**Figure S6.**
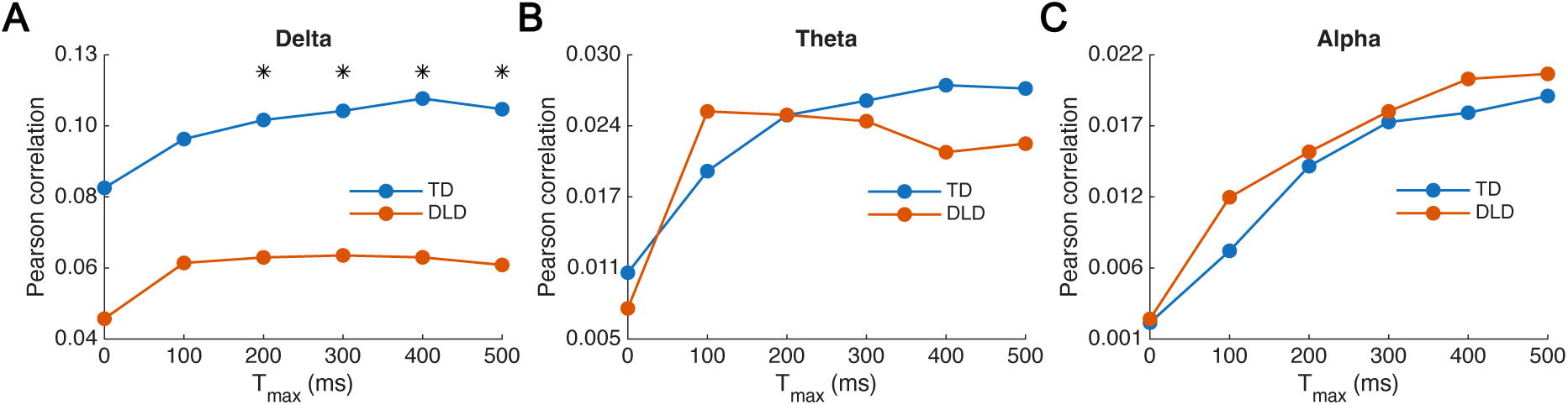
Within-child backward TRF reconstruction accuracy across delays (right temporal region). Stimulus reconstruction accuracy as a function of temporal delay (0–500 ms) for within-child backward TRF models restricted to the predefined right temporal electrode cluster. Panels show (A) delta, (B) theta, and (C) alpha bands. Values represent group-averaged decoding accuracy for TD (blue) and DLD (orange) children.

**Figure S7.**
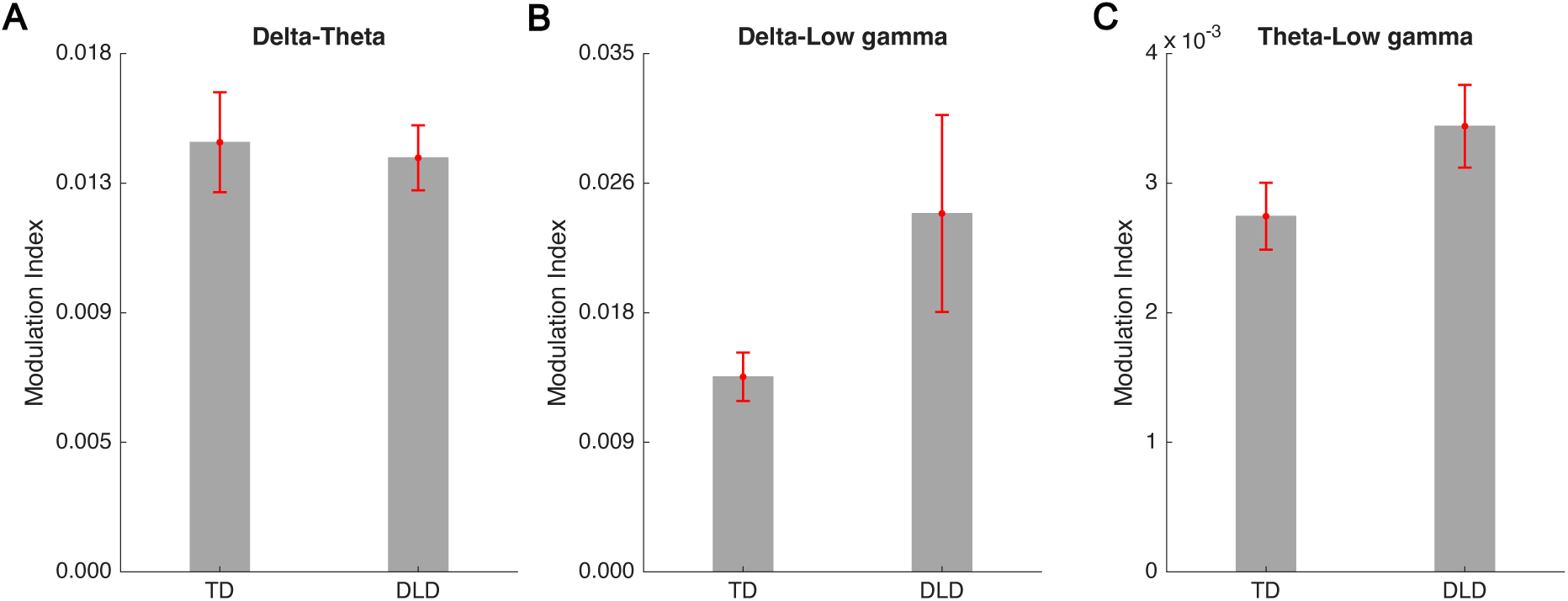
PAC measures by group (TD and DLD) and band. The modulation index was utilised as a measure for delta-theta (Panel A), delta-low gamma (Panel B), and theta-low gamma (Panel C) PAC. The error bars show the standard error of the mean.

**Figure S8.**
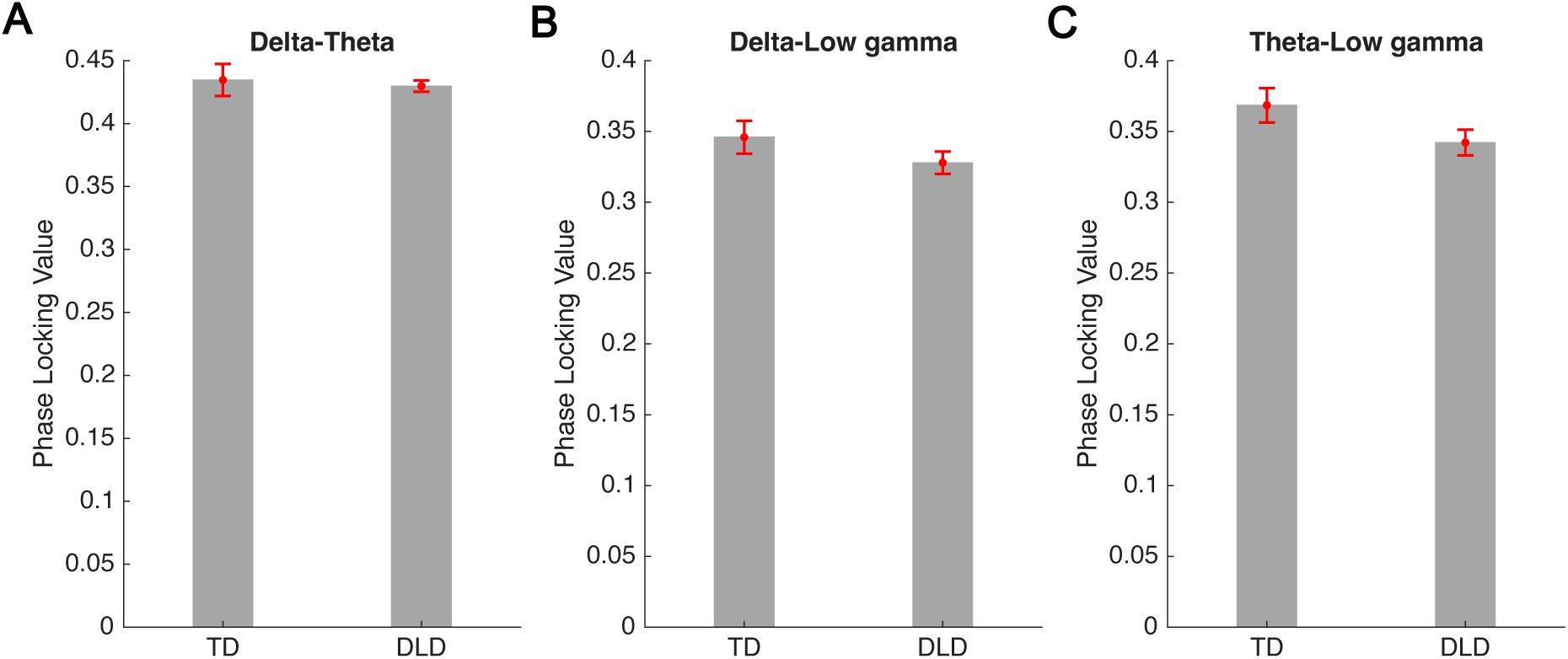
PPC measures by group (TD and DLD) and band. The phase locking value was utilised as a measure for delta-theta (Panel A), delta-low gamma (Panel B), and theta-low gamma (Panel C) PPC. The error bars show the standard error of the mean.

## Notes

### Competing Interest Statement

The authors have declared no competing interest.

### Summary of Updates

We have updated the Methods. The Introduction, Results, and Discussion sections have been revised accordingly.

## References

Araújo, J., Simons, B.D., Peter, V., Mandke, K., Kalashnikova, M., Macfarlane, A., Gabrielczyk, F., Wilson, A.M., Di Liberto, G.M., Burnham, D., & Goswami, U. (2024). Atypical low-frequency cortical encoding of speech identifies children with developmental dyslexia. Frontiers in Human Neuroscience, 18, 1403667. 10.3389/fnhum.2024.1403677

Asaridou, S.S., & Watkins, K.E. (2022). Neural basis of speech and language impairments in development: the case of developmental language disorder. (In K. Cohen Kadosh, Ed.; The Oxford Handbook of Developmental Cognitive Neuroscience). Oxford University Press. 10.1093/oxfordhb/9780198827474.013.19

Attaheri, A., Ní Choisdealbha, A., Di Liberto, G., Rocha, S., Brusini, P., Mead, N., Olawole-Scott, H., Boutris, P., Gibbon, S., Williams, I., Grey, C., Flanagan, S., M., & Goswami, U. (2022). Delta- and theta-band cortical tracking and phase-amplitude coupling to sung speech by infants. NeuroImage, 247, 118698. 10.1016/j.neuroimage.2021.118698

Attaheri, A., Ní Choisdealbha, A., Rocha, S., Brusini, P., Di Liberto, G., Mead, N., Olawole-Scott, H., Boutris, P., Gibbon, S., Williams, I., Grey Alfaro e Oliveira, M., Brough, C., Flanagan, S., M., & Goswami, U. (2024, in press). Infant low-frequency EEG cortical power, cortical tracking and phase-amplitude coupling predicts language a year later. PLoS One.

Beattie, R., & Manis, F. (2012). Rise time perception in children with reading and combined reading and language difficulties. Journal of Learning Disabilities, 46(3), 200–209. 10.1177/0022219412449421

Bedoin, N., Brisseau, L., Molinier, P., Roch, D., & Tillmann, B. (2016). Temporally Regular Musical Primes Facilitate Subsequent Syntax Processing in Children with Specific Language Impairment. Frontiers in Neuroscience, 10, 245. 10.3389/fnins.2016.00245

Benasich, A.A., Gou, Z., Choudhury, N., & Harris, K.D. (2008). Early cognitive and language skills are linked to resting frontal gamma power across the first three years. Behavioural Brain Research, 195, 215–222. https://doi.1016/j.bbr.2008.08.049

Bishop, D.V.M. (2013). Uncommon Understanding: Development and Disorders of Language Comprehension in Children. Psychology Press, 288, eBook ISBN9781315804699. 10.4324/9781315804699

Bishop, D., Snowling, M., Thompson, P., & Greenhalgh, T. (2017). Phase 2 of CATALISE: a multinational and multidisciplinary Delphi consensus study of problems with language development: Terminology. Journal of Child Psychology and Psychiatry, 58(10), 1068–1080. 10.1111/jcpp.12721

Cantiani, C., Riva, V., Piazza, C., Bettoni, R., Molteni, M., Choudhury, N., Marino, C., & Benasich, A.A., (2016). Auditory discrimination predicts linguistic outcome in Italian infants with and without familial risk for language learning impairment. Developmental Cognitive Neuroscience, 20, 23–34. 10.1016/j.dcn.2016.03.002

Cantiani, C., Ortiz-Mantilla, S., Riva, V., Piazza, C., Bettoni, R., Musaccia, G., Molteni, M., Marino, C., & Benasich, A. A., (2019). Reduced left-lateralized pattern of event-related EEG oscillations in infants at familial risk for language and learning impairment. Neuroimage: Clinical, 22, 101778. 10.1016/j.nicl.2019.101778

Choudhury, N., & Benasich, A.A. (2011). Maturation of auditory evoked potentials from 6 to 48 months: Prediction to 3- and 4-year language and cognitive abilities. Clinical Neurophysiology, 122, 320–338. 10.1016/j.clinph.2010.05.035

Corriveau, K.H., & Goswami, U. (2009). Rhythmic motor entrainment in children with speech and language impairments: Tapping to the beat. Cortex, 45, 119–130. 10.1016/j.cortex.2007.09.008

Corriveau, K., Pasquini, E., & Goswami, U. (2007). Basic auditory processing skills and specific language impairment: A new look at an old hypothesis. Journal of Speech, Language, and Hearing Research, 50, 647–666. 10.1044/1092-4388(2007/046)

Crosse, M.J., Di Liberto, G. M., Bednar, A., & Lalor, E.C. (2016). The multivariate temporal response function (mTRF) toolbox: a MATLAB toolbox for relating neural signals to continuous stimuli. Frontiers in Human Neuroscience, 10, 604. 10.3389/fnhum.2016.00604

Cumming, R., Wilson, A., & Goswami, U. (2015). Basic auditory processing and sensitivity to prosodic structure in children with specific language impairments: A new look at a perceptual hypothesis. Frontiers in Psychology, 6, 972. 10.3389/fpsyg.2015.00972

Daikoku, T., & Goswami, U. (2022). Hierarchical amplitude modulation structures and rhythm patterns: Comparing Western musical genres, song, and nature sounds to Babytalk. PLoS One, 17(10), e0275631. 10.1371/journal.pone.0275631

Destoky, F., Bertels, J., Niesen, M., Wens, V., Vander Ghinst, M., Leybaert, J., Lallier, M., Ince, R.A., Gross, J., De Tiège, X. and Bourguignon, M. (2020). Cortical tracking of speech in noise accounts for reading strategies in children. PLoS Biology, 18(8), p.e3000840. 10.1371/journal.pbio.3000840

Destoky, F., Bertels, J., Niesen, M., Wens, V., Vander Ghinst, M., Rovai, A., … & Bourguignon, M. (2022). The role of reading experience in atypical cortical tracking of speech and speech-in-noise in dyslexia. NeuroImage, 253, 119061. 10.1016/j.neuroimage.2022.119061

Di Liberto, G.M., Peter, V., Kalashnikova, M., Goswami, U., Burnham, D., & Lalor, E.C. (2018). Atypical cortical entrainment to speech in the right hemisphere underpins phonemic deficits in dyslexia. NeuroImage, 175, 70–79. 10.1016/j.neuroimage.2018.03.072

Donoghue, T., Haller, M., Peterson, E. J., Varma, P., Sebastian, P., Gao, R., … & Voytek, B. (2020). Parameterizing neural power spectra into periodic and aperiodic components. Nature neuroscience, 23(12), 1655–1665. 10.1038/s41593-020-00744-x

Fraser, J., Goswami, U., & Conti-Ramsden, G. (2010). Dyslexia and specific language impairment: The role of phonology and auditory processing. Scientific Studies of Reading, 14, 8–29. 10.1080/10888430903242068

Giraud, A-L., & Poeppel, D. (2012). Cortical oscillations and speech processing: Emerging computational principles and operations. Nature Neuroscience, 15, 511–517. 10.1038/nn.3063

Goswami, U. (2011). A temporal sampling framework for developmental dyslexia. Trends in Cognitive Sciences, 15(1), 3–10. 10.1016/j.tics.2010.10.001

Goswami, U. (2015). Sensory theories of developmental dyslexia: Three challenges for research. Nature Reviews Neuroscience, 16, 43–54. 10.1038/nrn3836

Goswami, U. (2022). Language acquisition and speech rhythm patterns: an auditory neuroscience perspective. Royal Society Open Science, 9(7), 211855. 10.1098/rsos.211855

Gross, J., Hoogenboom, N., Thut, G., Schyns, P., Panzeri, S., Belin, P., Garrod, S. (2013). Speech rhythms and multiplexed oscillatory sensory coding in the human brain. PLoS Biology, 11(12), e1001752. 10.1371/journal.pbio.1001752

Gou, Z., Choudhury, H., & Benasich, A.A. (2011). Resting frontal gamma power at 16, 24 and 36 months predicts individual differences in language and cognition at 4 and 5 years. Behavioural Brain Research, 220, 263–270. j.bbr.2011.01.048

Greenberg, S. (2006). A multi-tier framework for understanding spoken language. In S. Greenberg & W. Ainsworth (Eds.), Listening to speech: An auditory perspective. Lawrence Erlbaum Associates.

Greenberg, S., Carvey, H., Hitchcock, L., Chang, S., 2003. Temporal properties of spontaneous speech—a syllable-centric perspective. Journal of Phonetics, 31(3-4), 465–485. 10.1016/j.wocn.2003.09.005

Heim, S., Friedman, J.T., Keil, A., Benasich, A.A. (2011). Reduced Sensory Oscillatory Activity during Rapid Auditory Processing as a Correlate of Language-Learning Impairment. Journal of Neurolinguistics, 24(5), 539–555. 10.1016/j.jneuroling.2010.09.006

Helenius, P., Parviainen, T., Paetau, R., & Salmelin, R. (2009). Neural processing of spoken words in specific language impairment and dyslexia. Brain, 132(7), 1918–1927. 10.1093/brain/awp134

Helenius, P., Sivonen, P., Parviainen, T., Isoaho, P., Hannus, S., Kauppila, T., Salmelin, R., & Isotalo, L. (2014). Abnormal functioning of the left temporal lobe in language-impaired children. Brain and Language, 130, 11–18. 10.1016/j.bandl.2014.01.005

Hill, A. T., Clark, G. M., Bigelow, F. J., Lum, J. A., & Enticott, P. G. (2022). Periodic and aperiodic neural activity displays age-dependent changes across early-to-middle childhood. Developmental Cognitive Neuroscience, 54, 101076. 10.1016/j.dcn.2022.101076

Jessen, S., Fiedler, L., Münte, T., & Obleser, J. (2019). Quantifying the individual auditory and visual brain response in 7-month-old infants watching a brief cartoon movie. NeuroImage, 202, 116060. 10.1016/j.neuroimage.2019.116060

Kalashnikova, M., Peter, V., Di Liberto, G.M., Lalor, E.C., & Burnham, D. (2018). Infant-directed speech facilitates seven-month-old infants’ cortical tracking of speech. Scientific Reports, 8(1), 1–8. 10.1038/s41598-018-32150-6

Keshavarzi, M., Mandke, K., Macfarlane, A., Parvez, L., Gabrielczyk, F., Wilson, A., Flanagan, S. & Goswami, U. (2022a). Decoding of speech information using EEG in children with dyslexia: Less accurate low-frequency representations of speech, not “Noisy” representations. Brain & Language, 235, 105198. 10.1016/j.bandl.2022.105198

Keshavarzi, M., Mandke, K., Macfarlane, A., Parvez, L., Gabrielczyk F., Wilson, A., & Goswami, U. (2022b). Atypical Delta-band Phase Consistency and Atypical Preferred Phase in Children with Dyslexia during Neural Entrainment to Rhythmic Audio-Visual Speech. NeuroImage: Clinical, 35, 103054. 10.1016/j.nicl.2022.103054

Keshavarzi, M., Richards, S., Feltham, G., Parvez, L. & Goswami, U. (2024a). Neural processing of rhythmic speech by children with developmental language disorder (DLD): an EEG study. Imaging Neuroscience, 2, 1–20. 10.1162/imag_a_00382

Keshavarzi, M., Ní Choisdealbha, A., Attaheri, A., Rocha, S., Brusini, P., Gibbon, S., Boutris, P., Mead, N., Olawole-Scott, H., Ahmed, H., Flanagan, S., Mandke, K., & Goswami U. (2024b). Decoding speech information from EEG data with 4-, 7- and 11-month-old infants: Using convolutional neural network, mutual information-based and backward linear models. Journal of Neuroscience Methods, 403, 110036. 10.1016/j.jneumeth.2023.110036

Lachaux, J.P., Rodriguez, E., Martinerie, J., & Varela, F. J. (1999). Measuring phase synchrony in brain signals. Human brain mapping, 8(4), 194–208. 10.1002/(SICI)1097-0193(1999)8:4<194::AID-HBM4>3.0.CO;2-C

Lehongre, K., Morillon, B., Giraud, A.L., & Ramus, F. (2013). Impaired auditory sampling in dyslexia: further evidence from combined fMRI and EEG. Frontiers in human neuroscience, 7, 454. 10.3389/fnhum.2013.00454

Lehongre, K., Ramus, F., Villiermet, N., Schwartz, D., & Giraud, A.L. (2011). Altered low-gamma sampling in auditory cortex accounts for the three main facets of dyslexia. Neuron, 72(6), 1080–1090. 10.1016/j.neuron.2011.11.002

Leong, V., & Goswami, U. (2015). Acoustic-emergent phonology in the amplitude envelope of child-directed speech. Plos One, 10(12), e0144411. 10.1371/journal.pone.0144411

Leong, V., Kalashnikova, M., Burnham, D. & Goswami, U. (2017). The temporal modulation structure of infant-directed speech. Open Mind, 1 (2), 78–90. 10.1162/OPMI_a_00008

Lum, J.A.G., Clark, G.M., Bigelow, F.J., & Endicott, P.G. (2022). Resting state electroencephalography (EEG) correlates with children’s language skills: Evidence from sentence repetition. Brain & Language, 230, 105137. 10.1016/j.bandl.2022.105137

Mehler, J., Jusczyk, P., Lambertz, G., Halsted, N., Bertoncini, J., Amiel-Tison, C., 1988. A precursor of language acquisition in young infants. Cognition, 29(2), 143–178. 10.1016/0010-0277(88)90035-2.

Meng, X., Sun, C., Du, B., Liu, L., Zhang, Y., Dong, Q., … & Nan, Y. (2022). The development of brain rhythms at rest and its impact on vocabulary acquisition. Developmental Science, 25(2), e13157. 10.1111/desc.13157

Menn, K.H., Ward, E.K., Braukmann, R., van den Boomen, C., Buitelaar, J., Hunnius, S., et al. (2022). Neural Tracking in Infancy Predicts Language Development in Children with and without Family History of Autism. Neurobiology of Language, 3(3): 495–514. 10.1162/nol_a_00074

Molinaro, N., Lizarazu, M., Lallier, M., Bourguignon, M., & Carreiras, M. (2016). Out-of-synchrony speech entrainment in developmental dyslexia. Human brain mapping, 37(8), 2767–2783. 10.1002/hbm.23206

Nazzi, T., Bertoncini, J., Mehler, J., 1998. Language discrimination by newborns: toward an understanding of the role of rhythm. Journal of Experimental Psychology: Human Perception and Performance, 24(3), 756. 10.1037//0096-1523.24.3.756

Nora, A., Rinkinen, O., Renvall, H., Arkkila, E., Smolander, S., Laasonen, M., & Salmelin, R. (2024). Impaired cortical tracking of speech in children with developmental language disorder. Journal of Neuroscience, 44(22). 10.1523/JNEUROSCI.2048-23.2024

Ortiz Barajas, M.C., Guevara, R., & Gervain, J. (2021). The origins and development of speech envelope tracking during the first months of life. Developmental Cognitive Neuroscience, 48, 100915. 10.1016/j.dcn.2021.100915

Parker, A.J., Woodhead, Z.V.J., Carey, D.P., Groen, M.A., Gutierrez-Sigut, E., Hodgson, Hudson, J., Karlsson, E.M., MacSweeney, M., Payne, H., Simpson, N., Thompson, P.A., Watkins, K.E., Egan, C., Grant, J.H., Harte, S., Hudson, B.T., Sablik, M., Badcock, N.A., & Bishop, D.V.M. (2022). Inconsistent language lateralisation: Testing the dissociable language laterality hypothesis using behaviour and lateralised cerebral blood flow. Cortex, 154, 105–134. 10.1016/j.cortex.2022.05.013

Petri, S., Vitali, H., Campus, C., Riberto, M., & Gori, M. (2025). Exploring neurodevelopment through oscillatory and aperiodic EEG activity: methodological and clinical consideration. Frontiers in Human Neuroscience, 19, 1641840. 10.3389/fnhum.2025.1641840

Power, A.J., Colling, L.C., Mead, N., Barnes, L., & Goswami, U. (2016). Neural encoding of the speech envelope by children with developmental dyslexia. Brain & Language, 160, 1–10. 10.1016/j.bandl.2016.06.006

Power, A.J., Mead, N., Barnes, L., & Goswami, U. (2013). Neural entrainment to rhythmic speech in children with developmental dyslexia. Frontiers in Human Neuroscience, 7, 777. 10.3389/fnhum.2013.00777

Przybylski, L., Bedoin, N., Krifi-Papoz, S., Herbillon, V., Roch, D., & Léculier, L., Kotz, S.A., & Tillmann, B. (2013). Rhythmic auditory stimulation influences syntactic processing in children with developmental language disorders. Neuropsychology, 27(1), 121–131. 10.1037/a0031277

Richards, S., & Goswami, U. (2015). Auditory processing in specific language impairment (SLI): Relations with the perception of lexical and phrasal stress. *Journal of Speech*, Language, and Hearing Research, 58(4), 1292–1305. 10.1044/2015_jslhr-l-13-0306

Richards, S., & Goswami, U. (2019). Impaired recognition of metrical and syntactic boundaries in children with developmental language disorders. Brain Sciences, 9(2), 33. 10.3390/brainsci9020033

Sallat, S., & Jentschke, S. (2015). Music perception influences language acquisition: Melodic and rhythmic-melodic perception in children with specific language impairment. Behavioural Neurology, 2015, 606470. 10.1155/2015/606470

Tort, A.B.L., Kramer, M.A., Thorn, C., & Kopell, N.J. (2008). Dynamic cross-frequency couplings of local field potential oscillations in rat striatum and hippocampus during performance of a T-maze task. Proceedings of the National Academy of Sciences, 105(51) 20517–20522. 10.1073/pnas.081052410

Wechsler, D. (2016). Wechsler Intelligence Scale for Children, 4th Edn. London, UK: Pearson Assessment.

Welch, P. (1967). The use of fast Fourier transform for the estimation of power spectra: a method based on time averaging over short, modified periodograms. IEEE Transactions on audio and electroacoustics, 15(2), 70–73.

Wiig, E.H., Semel, E., Secord, W.A. (2017). Clinical Evaluation of Language Fundamentals - Fifth Edition. Pearson.

